# Divergent Behavioral and Circuit-Level Adaptations to Acute and Chronic Gastric Electrical Stimulation

**DOI:** 10.64898/2026.05.20.726694

**Authors:** Maryana Daood, Rumail Memon, Spandan Rout, Bassant Elgengihy, Alec Nossa, Robert Feldman, Mahmoud Elbeh, Sadaf Usmani, Khalil B Ramadi

**Author notes:** College of Medicine, Department of Physiology and Aging, University of Florida, Gainesville, FL.

## Abstract

Anxiety disorders are highly prevalent and often refractory to existing treatments, motivating the development of alternative neuromodulatory strategies. Peripheral bioelectronic approaches targeting the gut–brain axis such as vagus nerve stimulation (VNS) demonstrate that modulation of visceral afferent pathways can influence central emotional circuits. Gastric electrical stimulation (GES) is a clinically established therapy for gastrointestinal motility disorders. While the stomach is densely innervated by vagal afferents, the effect of GES on anxiety-related behavior has not been systematically examined. We sought to identify neural pathways engaged by GES and effects of continuous chronic GES on behavior. To do this, we developed a gastric stimulation platform for rodents, a fully implantable, untethered system enabling chronic neuromodulation in freely moving rats. We combined this with cross-species whole-brain activity mapping in mice to interrogate circuit-level mechanisms. Using open field and elevated plus mazes, together with machine-learning-based behavioral tracking and multivariate modeling, we show that acute GES induces a robust, context-dependent anxiogenic phenotype characterized by reduced exploration and increased freezing, particularly in novel open-field environments. In contrast, chronic GES produces a divergent post-stimulation behavioral profile marked by enhanced exploratory behavior relative to acutely stimulated animals, indicating temporally dynamic reorganization of anxiety-related behavior. Principal component analysis and hierarchical clustering further revealed that stimulation reshapes the multivariate structure of behavioral features rather than shifting animals along a single anxiety continuum. Whole-brain c-Fos mapping revealed anatomically distributed modulation of limbic–cortical networks following gastric stimulation, including suppression of ventral medial entorhinal cortex, excitation of nucleus tractus solitarii and heterogeneous recruitment of amygdalar and hippocampal subregions. These circuit-level patterns align with the behavioral dissociation between contextual exploration and explicit threat avoidance, providing convergent cross-species evidence that gastric stimulation engages distributed anxiety-related networks. Together, these findings establish the first freely moving behavioral model of chronic gastric neuromodulation, demonstrate temporally dynamic and context-sensitive effects on anxiety-like behavior, and provide systems-level validation that the stomach can serve as a viable peripheral access point for modulating central emotional circuits.

## INTRODUCTION

Anxiety disorders are the most common and debilitating mental illnesses worldwide (1). Despite the availability of psychotherapeutic and pharmacological treatments, a substantial proportion of patients remain resistant to therapy, and only about one-quarter of those who seek professional help receive adequate treatment (2). This therapeutic gap has spurred interest in alternative approaches, including neuromodulation techniques, to modulate brain circuitry and alleviate anxiety. One promising avenue leverages the bidirectional communication along the gut–brain axis, whereby signals from the gastrointestinal (GI) tract influence central nervous system function and emotional states (3). This gut-brain communications is based on a complex system, including sympathetic, endocrine, immune, microbiome and humoral links in addition to the vagus nerve being a key conduit, with the majority of its fibers carrying interoceptive information from visceral organs to the brainstem (3–5). Notably, vagal afferent signaling has been implicated in the regulation of mood and affect, suggesting that peripheral interventions targeting the vagus or its end-organs could impact anxiety and fear processing (6, 7). Indeed, patients with functional GI disorders commonly present with anxiety comorbidities; for example anxiety disorders are diagnosed in about 38% of functional dyspepsia patients, versus 4% in general population (8), underscoring the intimate connection between gut physiology and emotional behavior.

Vagus nerve stimulation (VNS) is an established neuromodulation therapy that capitalizes on this gut–brain neural pathway, see Fig. 1B. Electrical stimulation of the cervical vagus nerve has been used clinically for refractory epilepsy and depression (9–11), and accumulating evidence points to its efficacy in anxiety-related conditions (6, 12–15). Preclinical studies in rodents show that certain parameters of VNS produce robust anxiolytic effects. For example, in vivo experiments have demonstrated that VNS at moderate intensities can significantly reduce anxiety-like behavior in standard paradigms such as the elevated plus maze and open field maze (6, 13). Correspondingly, VNS-treated rats exhibit neural activation changes in brainstem and limbic regions; mid-intensity VNS selectively engages nucleus tractus solitarius (NTS) neurons, including noradrenergic cells, and suppresses locus coeruleus activity, correlating with diminished anxiety-like responses (6). These findings align with a broader body of work indicating that vagal afferent stimulation can modulate neurotransmitter systems involved in stress and fear (i.e. noradrenaline and GABA in the limbic circuitry) thereby altering emotional behavior (7). Hence, VNS provides a precedent that peripheral neuromodulation of vagal pathways can induce anxiolytic effects, motivating exploration of other vagal-associated targets along the gut–brain axis for treating anxiety.

**Figure 1.**
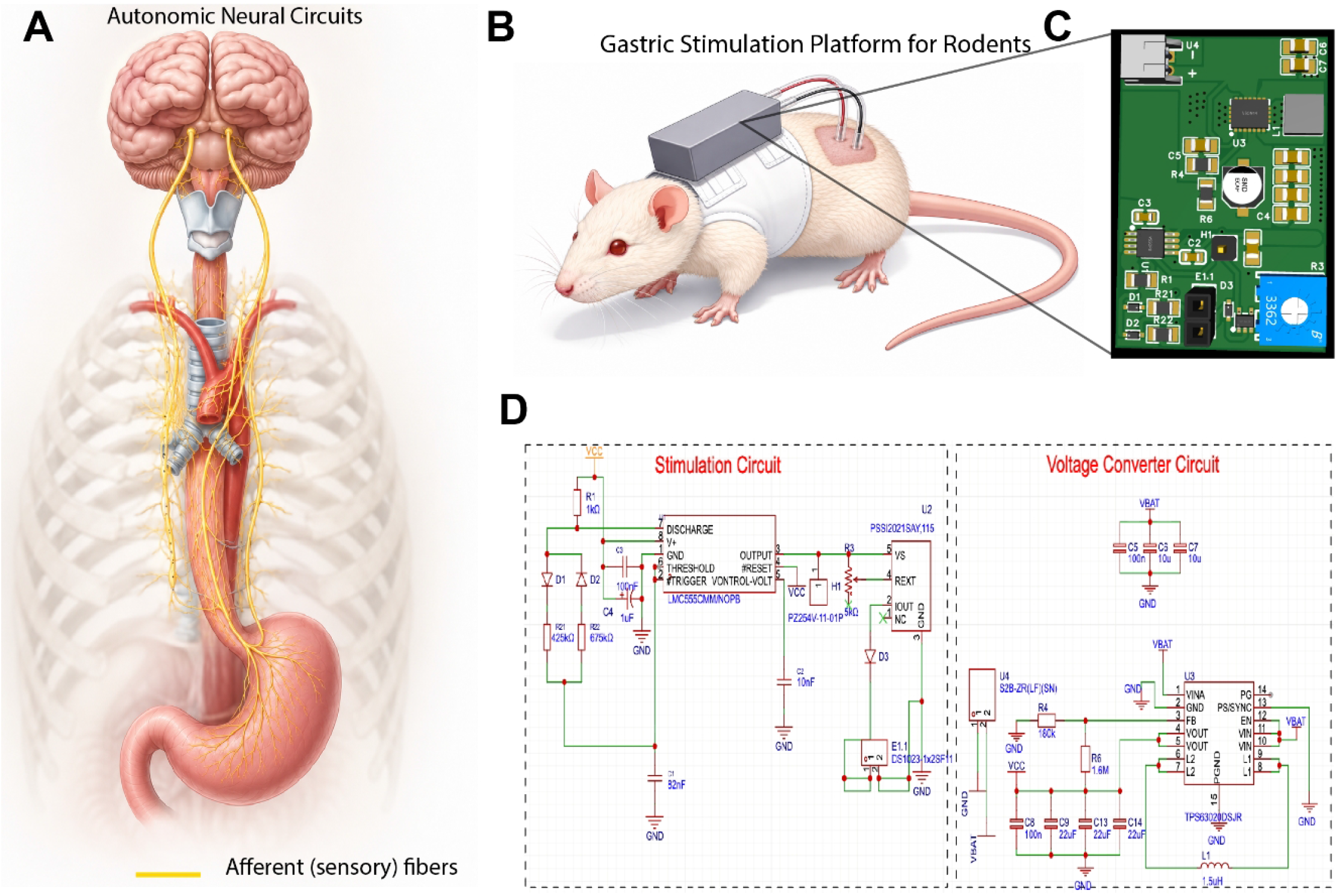
Summary of the untethered gastric electrical stimulation (GES) system. (A) Schematic of the gut-brain axis (B) Schematic of the fully implantable, dorsal-mounted backpack system worn by a rat during free behavior. (C) Custom microstimulator printed circuit board (PCB). (D) Circuit schematic illustrating a 555 timer–based pulse generator driving a constant-current output stage. Stimulation parameters: monophasic, frequency = 14 Hz, pulse width = 0.3 ms, current amplitude = 0.5 mA. Acute stimulation consisted of a single 30-min session. Chronic stimulation consisted of two 48-h stimulation blocks separated by 24-h.

Gastric electrical stimulation (GES) is a neuromodulatory technique that delivers mild electrical pulses to the stomach, an organ richly innervated by vagal afferents (4). Clinically, GES has been applied for decades to manage refractory gastroparesis and other GI motility disorders (16–18). The mechanism of GES is thought to involve modulating enteric neural circuits and vagal nerve endings in the stomach wall, thereby influencing gastric myoelectrical activity as well as afferent signaling to the brain (19, 20). Recent evidence supports the notion that stomach-targeted stimulation can have far-reaching neural effects: for instance, Cao and colleagues showed that electrical stimulation of the rodent stomach elicited rapid and widespread fMRI activity changes in multiple brain regions, including somatosensory and cingulate cortices (21). Such findings suggest GES can engage central autonomic and emotional networks via the vagus nerve. Given that approximately 80% of vagal fibers are sensory afferents transmitting information from gut to brain (3), stimulating the stomach might tap into similar pathways, potentially modulating mood and anxiety-related circuits. This raises the possibility that therapeutic gastric stimulation could be harnessed to favorably influence mental health, akin to how VNS is used in neuropsychiatric disorders.

Novel bioelectronic medicine techniques utilize ingestible and gastrointestinal electroceuticals that interface directly with the gut to modulate systemic physiology (22–28). While such technologies are being developed to regulate metabolism and motility, their potential to influence emotional and behavioral states has not been systematically investigated. To date, no study has directly evaluated whether targeted gastrointestinal electrical stimulation alters anxiety-related behavior in freely moving animals. Thus, whether the stomach can serve as an entry point for behavioral neuromodulation remains an open and clinically relevant question.

Prior GES research has predominantly focused on gastrointestinal symptoms and motility outcomes, and no study to has directly evaluated whether chronic GES can modulate anxiety-like behaviors. A major technical hurdle has been the lack of a suitable preclinical model for delivering GES during complex behavioral assays. Traditional experimental setups often involve tethered stimulation equipment or acute anesthetized preparations (6, 13, 14, 29), which are not conducive to observing naturalistic behavior in freely moving animals. Here, we address this gap by developing a novel untethered GES platform for rodents, enabling chronic stimulation in free-moving rats during behavioral testing. Our backpack-mounted stimulator system allowed us to implant electrodes in the stomach and administer GES without restraint, thus preserving normal exploratory and anxiety-related behaviors even during the stimulation. We examined both acute and chronic gastric stimulation and found that GES produces temporally dynamic and context-dependent effects on anxiety-like behavior. We find that acute stimulation increases anxiety-like behavior in the open field maze, whereas chronic stimulation produces a divergent post-stimulation exploratory phenotype. These findings indicate that gastric neuromodulation does not exert a uniform behavioral effect but instead reorganizes anxiety-related behavior in a time-dependent manner. To further resolve the neural circuits engaged by gastric stimulation, we complemented our rat behavioral studies with whole-brain activity mapping in mice. This cross-species approach allowed circuit-level interrogation of gut–brain activation patterns not yet practical at comparable resolution and throughput in the larger rat brain, providing anatomical validation of the pathways potentially underlying the observed behavioral effects. Together, this work establishes the first freely moving behavioral model of gastric electrical stimulation, situates GI neuromodulation within the expanding framework of ingestible bioelectronic therapeutics, and provides both behavioral and whole-brain circuit evidence that the stomach can serve as a viable access point for modulating central anxiety-related neural networks.

## RESULTS

### Development and validation

We developed an untethered GES system that could be chronically implanted and functional for up to 14 days. The system consists of a backpack mounted on a jacket (Fig.1B). The custom microstimulator board (Fig.1C) with the custom stimulation system (Fig.1D) measured approximately 2.5 cm x 2 cm and housed a rechargeable battery and a controller. Ultra fine lead Platinum iridium wires (.003” diameter, ~ 76 μm) were used as stimulating electrodes, threaded through the stomach wall and secured onto the gastric serosa (Fig. 2A-D, S1D). We removed insulating coating from a small section (0.5cm) of electrode wires to ensure adequate gastric mucosal electrical contact (Fig. S1C).

**Figure 2.**
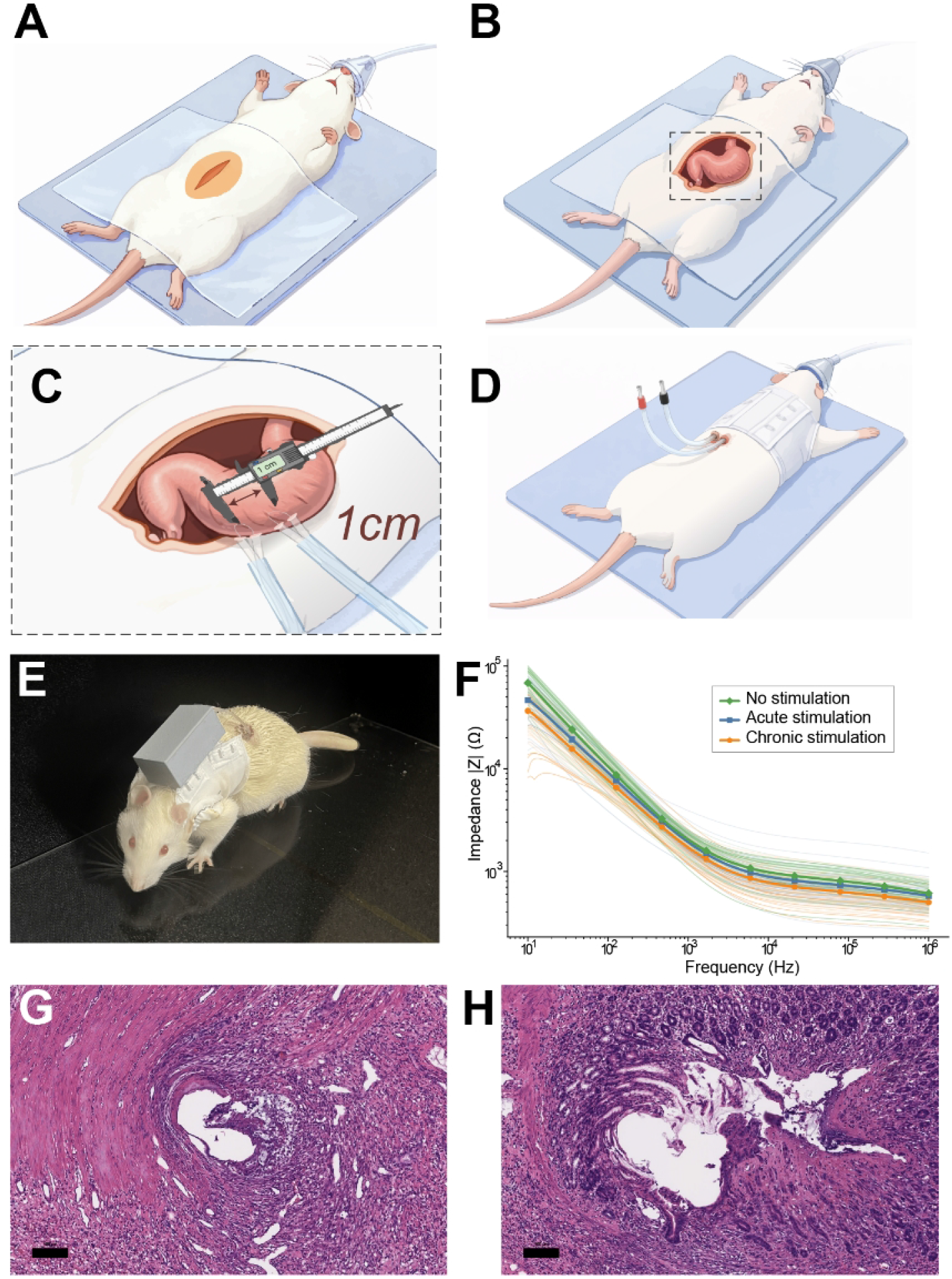
Development, implantation, and validation of the untethered gastric electrical stimulation (GES) system. (A–D) Stepwise schematic of electrode implantation: (A) lateral abdominal incision; (B) stomach exteriorization; (C) implantation of two 14.5-inch platinum–iridium wires (0.003” diameter) into the greater curvature (~1 cm spacing); (D) subcutaneous routing of leads to a dorsal exit site. (E) Representative image of a rat during behavioral testing while wearing the dorsal stimulator. (F) Electrode impedance spectra across groups (Acute GES n = 3; Chronic GES n = 4; No stimulation n = 5). Impedance spectra were acquired across 125 logarithmically spaced frequencies ranging from 10 Hz to 100 kHz. Spectra were compared against a reference range obtained when electrodes were in contact with the mucosal surface. Group differences assessed using two-sided cluster-based permutation testing across frequencies; no clusters survived correction (cluster-level α = 0.05). (G) Representative hematoxylin and eosin (H&E) staining from stomach tissue implanted with 0.003” electrodes (n = 5, 9 sections analyzed). (H) Representative H&E staining from 0.005” electrodes (n = 5, 9 sections analyzed). Histological assessment was qualitative; no statistical comparison was performed.

Impedance monitoring demonstrated a stable electrode-tissue interface throughout the experiment. Impedance values for each rat stayed consistent within a narrow range (± 1SD band) during the entire protocol period (see Fig. S1H for sample individual rat plots). There were no significant differences in impedance trends between the groups (acute GES, chronic GES, no stimulation; Fig. 2F). A cluster-based permutation analysis of the impedance time series revealed no group effect (corrected *p* ≥ 0.05 for all pairwise group comparisons), indicating that all groups maintained similar electrode integrity, and the GES did not degrade electrode performance.

Postmortem inspection confirmed correct electrode placement and negligible fibrotic tissue encapsulation around the leads, indicating successful long-term implantation and biocompatibility (Fig. S1E). Evaluation of the histology stomach of the stimulation sites of .003” electrodes (Fig. 2G) by a board-certified pathologist showed no significant morphological changes, bleeding, or tissue damage in comparison to .005” electrodes (Fig. 2H), supporting the safety of our untethered gastric stimulation model.

### Acute and chronic behavioral effects of GES

#### Baseline (pre-stimulation) group differences

One-way ANOVAs revealed no significant baseline differences among the three groups (n = 6, 5, 6 for Groups 1,3, and 4, respectively) on any measured behavioral metric in either the EPM or OFM tasks. For example, the % time spent on open arms in the EPM during the pre-stimulation epoch was low and comparable across groups (Group 1: 9.745% vs Group 3: 13.29% vs Group 4: 8.2%, F < 1, p > .05), see Fig.S2A. Similarly, baseline OFM anxiety-related measures (e.g. % time in center, freezing duration) did not differ significantly between groups (all p > .05), see Fig.S2B. These results indicate that the groups were behaviorally equivalent at baseline, before any stimulation.

#### During stimulation group differences

During the stimulation epoch, significant between-group differences in one-way ANOVAs emerged on several measures. In the OFM, the groups differed in velocity (cm/s)(F (3, 15) = 3.347, p = .047, η^2^ = .381). Acute stim group (Group 1) had significantly decreased velocity than the control group (Group 4) (p = .036), Fig 3A. Furthermore, significant group effect emerged in grooming rate (F (3, 14) = 4.838, p = .016, η^2^ = .380). Acute stim group (Group 1) had significantly decreased grooming rate than the sham surgery group (Group 3) (p = .034), Fig 3B. Additionally, significant group effect emerged in % freeze time (F (3, 15) = 5.112, p = .012, η^2^ = .456). Acute stim group (Group 1) had significantly increased % freeze time than the sham surgery group (Group 3) (p = .009) and the control (Group 4) (p = .032), Fig. 3C. Chronic stimulation (Group 2) showed a similar trend in most measures (intermediate between acute GES and controls), but differences were not statistically significant after correction (all p > .05). Focusing on the acute GES group (Group 1), we further segmented the 30 minutes stimulation (Stim1, Stim2, Stim3) to examine whether this decreased exploration is a function of time spent in the maze or a cumulative effect of stimulation. Linear mixed-effects models with epoch (*Pre-stim, During-stim Stim1, Stim2, Stim3, Post-stim*) and repetitions (*Repetition* 1, 2 and 3) as a within-subject factor and rat as random intercept revealed a main effect of epoch for velocity (F (1.176, 5.879) = 13.24, p = .01). Post hoc tests revealed that velocity has decreased significantly from a *Pre-stim* mean to *Stim3* on the first *repetition* (p = .027, mean diff.= 1.072 ± .229, n = 6) and on the second *repetition* (p = .019, mean diff.= 1.113 ± .149, n = 4) and on the third *repetition* (p < .001, mean diff.= 1.143 ± .027, n = 3; Fig. 3G). No significant ANOVA’s emerged in the EPM (all p > .05)

**Figure 3.**
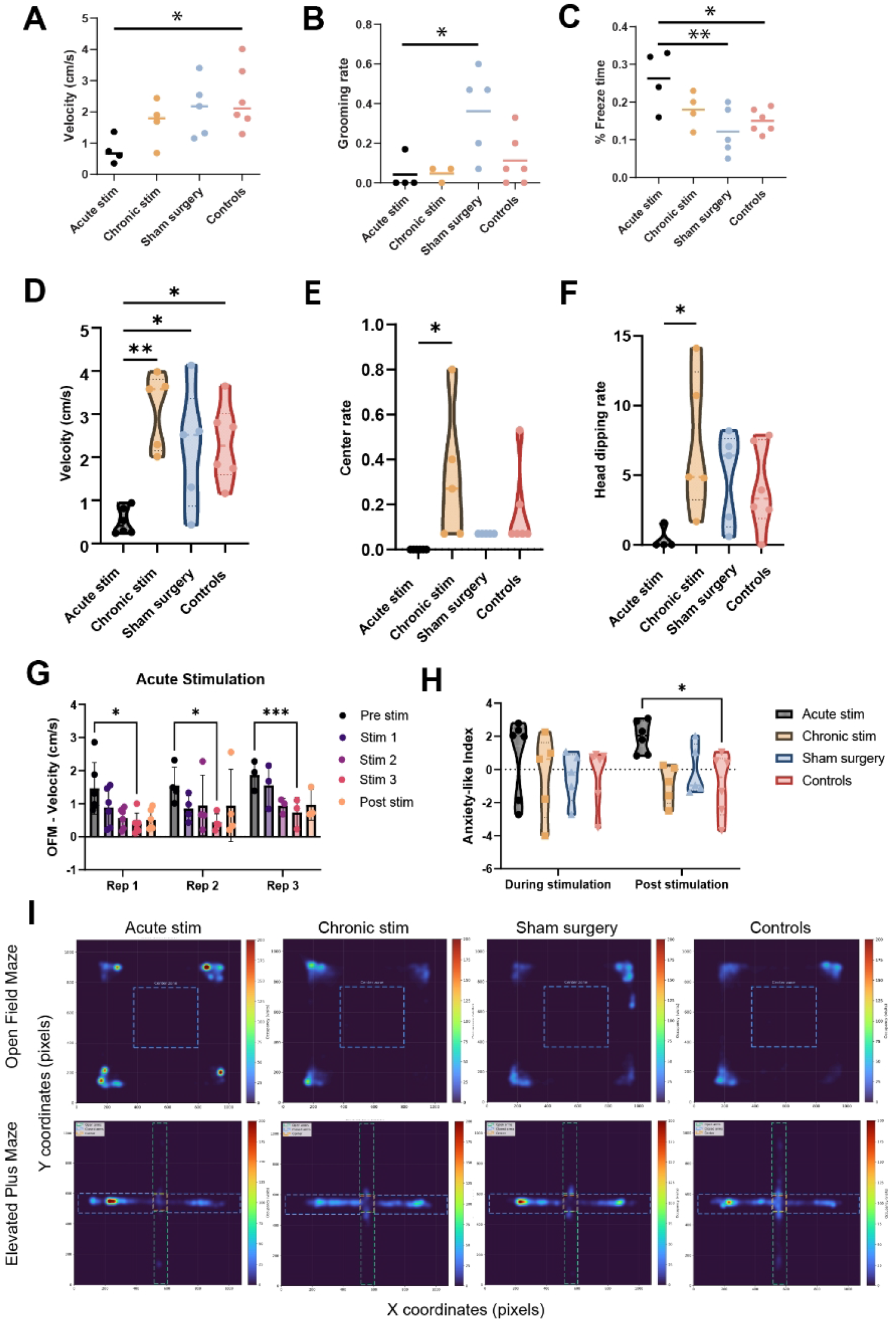
Behavioral effects of acute and chronic gastric stimulation. Group sizes: Acute GES n = 4; Chronic GES n = 4; Sham n = 5; No-surgery control n = 6. All statistical tests were two-sided. (A) The groups differed in velocity (cm/s) in the Open Field Maze (OFM; F (3, 15) = 3.347, p = .047, η^2^ = .381). Acute stim group (Group 1) had significantly decreased velocity than the control group (Group 4) (p = .036). (B) Significant group effect emerged in grooming rate in the OFM (F (3, 14) = 4.838, p = .016, η^2^ = .380). Acute stim group (Group 1) had significantly decreased grooming rate than the sham surgery group (Group 3) (p = .034). (C) Significant group effect emerged in % freeze time in the OFM (F (3, 15) = 5.112, p = .012, η^2^ = .456). Acute stim group (Group 1) had significantly increased % freeze time than the sham surgery group (Group 3) (p = .009) and the control (Group 4) (p = .032). (D) Significant group effect emerged in velocity (cm/s) in the OFM (F (3, 18) = 7.736, p = .001, η^2^ = .619). Acute stim group (Group 1) had significantly decreased velocity than the chronic stim group (Group 2) (p = .001), and the sham surgery group (Group 3) (p = .036) and the control group (Group 4) (p = .017). (E) Significant group effect emerged in center rate in OFM (F (3, 18) = 3.500, p = .036, η^2^ = .368). Acute stim group (Group 1) had significantly decreased center rate than the chronic stim group (Group 2) (p = .029). (F) Group effect trend emerged in head dipping rate in EPM (F (3, 16) = 2.885, p = .068, η^2^ = .350). Acute stim group (Group 1) had significantly decreased head dipping rate than the chronic stim group (Group 2) (p = .045). (G) OFM velocity segmented into time bins across the 30-min acute stimulation session (first 10 minutes, second 10 minutes, third 10 minutes). Linear mixed-effects models with epoch (*Pre-stim, Stim1, Stim2, Stim3, Post-stim*) and repetitions (*Repetition* 1, 2 and 3) as a within-subject factor and rat as random intercept revealed a main effect of epoch for velocity (F (1.176, 5.879) = 13.24, p = .01). Post hoc Tukey tests revealed that velocity has decreased significantly from a *Pre-stim* mean to *Stim3* on the first *repetition* (p = .027, mean diff.= 1.072 ± .229, n = 6) and on the second *repetition* (p = .019, mean diff.= 1.113 ± .149, n = 4) and on the third *repetition* (p < .001, mean diff.= 1.143 ± .027, n = 3). (H) Principal component–derived AnxietyIndex (PC1) during OFM testing. Two-way ANOVA with *Group* (four levels) as between-subject and *Epoch* (during vs. post-stimulation) within-subject factors revealed a main effect trend of *Group* on *AnxietyIndex* (F(3, 18) = 2.677, *p* = 0.078) with the acute stim group (Group 1) being the highest on the *AnxietyIndex* score. Post-hoc Tukey tests indicated that Group 1 (Acute GES) exhibited significantly higher *AnxietyIndex* scores than the controls (Group 4) (p = 0.041). (I) Occupancy heatmaps (XY position density) in OFM and EPM for each group. Data are shown as mean ± SD. *p < 0.05 (FDR corrected).

#### Post stimulation group differences

Post stimulation epoch, significant between-group differences in one-way ANOVAs emerged on several measures. In the OFM, the groups differed in velocity (cm/s)(F (3, 18) = 7.736, p = .001, η^2^ = .619). Acute stim group (Group 1) had significantly decreased velocity than the chronic stim group (Group 2) (p = .001), and the sham surgery group (Group 3) (p = .036) and the control group (Group 4) (p = .017), Fig 3D. Furthermore, significant group effect emerged in center rate (F (3, 18) = 3.500, p = .036, η^2^ = .368). Acute stim group (Group 1) had significantly decreased center rate than the chronic stim group (Group 2) (p = .029), Fig 3E. Additionally, a group effect trend emerged in head dipping rate in EPM (F (3, 16) = 2.885, p = .068, η^2^ = .350). Acute stim group (Group 1) had significantly decreased head dipping rate than the chronic stim group (Group 2) (p = .045), Fig. 3F.

Overall heatmaps show that across all epochs, acute stimulation group (Group 1) spent more time in corner coordinates compared to the other groups, indicating decreased velocity and increased freezing (Fig. 3I).

### Multivariate behavioral effects

To capture a latent dimension of anxiety-like behavior, we conducted principal component analysis (PCA) on standardized behavioral metrics from the OFM and EPM tasks. In the OFM, PCA on 11 behavioral variables revealed that the first principal component (PC1) accounted for 31.9% of the total variance. For the EPM, PC1 explained 39.8% of the variance across eight anxiety-related features, Fig S3A-B. In both tasks, PC1 reflected a behavioral pattern consistent with elevated anxiety marked by increased freezing and reduced exploratory activity. The direction of PC1 was inverted such that higher scores corresponded to greater anxiety-like behavior; this resulting metric (*AnxietyIndex)* and used as the primary dependent variable for subsequent analyses.

We next examined the effects of GES on *AnxietyIndex* scores using a two-way ANOVA with *Group* (four levels) as between-subject and *Epoch* (during vs. post-stimulation) within-subject factors was conducted. In the OFM, the ANOVA revealed a main effect trend of *Group* on *AnxietyIndex* (F(3, 18) = 2.677, *p* = 0.078) with the acute stim group (Group 1) being the highest on the *AnxietyIndex* score. Post-hoc Tukey tests indicated that Group 1 (Acute GES) exhibited significantly higher *AnxietyIndex* scores than the controls (Group 4) (p = 0.041). All other pairwise group comparisons were not statistically significant (all *p* > 0.05), Fig 3H.

In the EPM task, the same two-way ANOVA showed no significant effects. Tukey-adjusted post-hoc comparisons confirmed no significant group differences in *AnxietyIndex* during or after stimulation (all *p* > 0.5).

We further conducted Pearson correlation analyses to examine the interrelationships among behavioral metrics collected from OFM and EPM tasks during and post stimulation to explore shared variance across measures. Several statistically significant FDR-corrected correlations emerged (p < 0.05), Fig. 4A. A few notable significant Pearson correlation noted; Within OFM, velocity post-stimulation showed a positive correlation with grooming rate post-stimulation (r = .680, p = .035, Fig. 4B), center entry rate post-stimulation (r = 0.630, p = 0.035, Fig. 4C). Within EPM, head dipping rate post-stimulation showed a positive correlation with open arms entries rate post-stimulation (r = 0.729, p = 0.035, Fig. 4D) and percentage of time spent in the open armspost-stimulation (r=0.762, p=0.035, Fig.4E) and open arms index (i.e 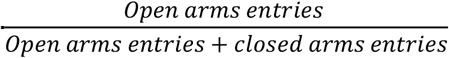) post-stimulation (r = 0.693, p = 0.035, Fig. 4F). Cross mazes, velocity post-stimulation in OFM was positively correlated with head dipping rate post-stimulation in EPM (r = 0.777, p = 0.035, Fig. 4G). Cross epochs, velocity during-stimulation in OFM was positively correlated with open arms entry rate post-stimulation in EPM (r = 0.730, p = 0.035, Fig. 4H) and finally SAP rate in EPM was positively correlated between during-stimulation and post-stimulation (r = .95, p = .035, Fig 4I).

**Figure 4.**
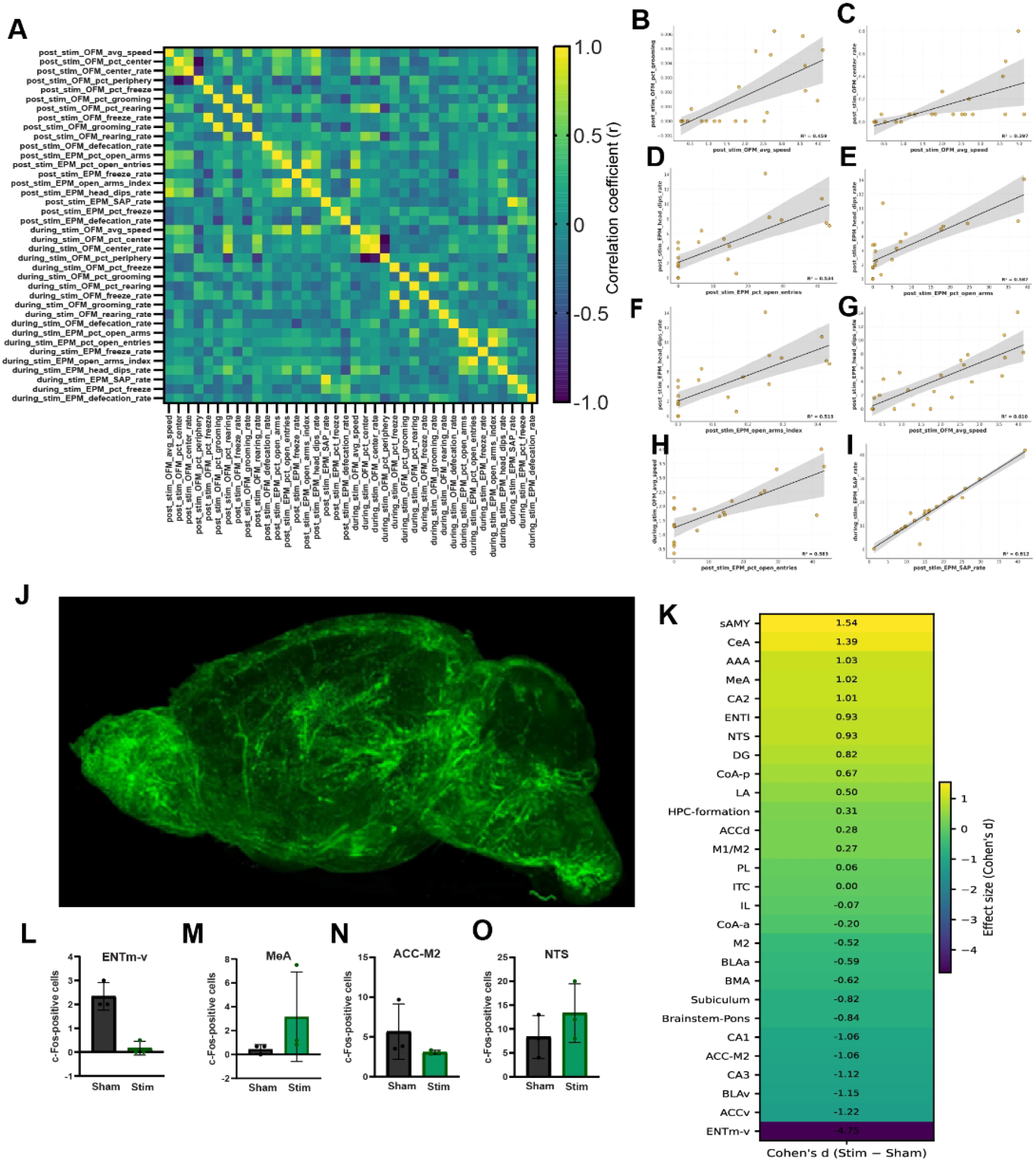
Multivariate behavioral structure and whole-brain activity mapping following gastric stimulation. Behavioral correlations computed across all animals (n = 18 total). Pearson correlation coefficients two-tailed were computed. FDR correction (Benjamini–Hochberg, q = 0.05) was applied across all matrix comparisons. (A) Correlation matrix of OFM and EPM variables across during- and post-stimulation epochs. (B–I) Representative significant Pearson correlation scatter plots with best-fit linear regression lines. (B) OFM velocity vs grooming rate (post); (C) OFM velocity vs center entries (post); (D–F) EPM head-dipping vs open-arm measures (post); (G) Cross-maze correlation: OFM velocity vs EPM head-dipping (post); (H) OFM velocity (during) vs EPM open-arm entries (post); (I) EPM stretch-attend posture rate (during vs post). (J) Whole-brain cFos immunolabeling following acute GES (Acute GES n = 3; Sham n = 3). Brains processed using CLARITY-based tissue clearing and immunostaining for cFos. Imaging performed using CLARITY-optimized light-sheet microscope with pixel resolution of 1.4 × 1.4 µm^2^ (x-y dimensions) and 5 µm z-step using a 4×/0.28 NA objective. (K) ROI-level effect size heatmap (Cohen’s d) illustrating heterogeneous modulation across limbic–cortical regions. (L–O) Representative ROIs: ventral medial entorhinal cortex (ENTm-v), medial amygdala (MeA), anterior cingulate cortex–secondary motor cortex border (ACC–M2), nucleus tractus solitarii (NTS). Cell detection was performed using automated threshold-based segmentation in cleared volumes. cFos+ cell counts were normalized to ROI volume (cells/mm^3^). Brains were registered to the Allen Rat Brain Atlas. Statistical comparison for each ROI used two-sided permutation testing (10,000 permutations). FDR correction (Benjamini–Hochberg, q = 0.05) was applied across all ROIs. No ROI survived FDR correction (all q ≥ .8). Data are presented as effect size (Cohen’s d). Exact p and q values are provided in Supplementary Table S4.

Additionally, we compared the hierarchical organization of behavioral variables across the four experimental groups using two independent dendrogram-comparison metrics: the cophenetic correlation coefficient (CCC) and Baker’s Gamma (Fig S3C). Both metrics quantify the degree to which two trees preserve similar clustering arrangements. Across all pairwise group comparisons, CCC values were extremely low (0.059–0.157) and Baker’s Gamma coefficients were close to zero (−0.008–0.077), indicating negligible similarity in variable clustering patterns between groups. These results demonstrate that the multivariate behavioral structure differs substantially across stimulation conditions, with no evidence of shared cluster architecture. Thus, each group exhibits a distinct pattern of relationships between behavioral measurements, suggesting that stimulation protocols reorganize behavioral features in different ways.

### Neural effects of GES

Whole-brain CLARITY imaging revealed distributed recruitment of cFos-positive neurons following stimulation, spanning limbic, frontal, and parahippocampal regions (Fig. 4J). Activated neurons were spatially organized along a medial cortical–limbic axis. Quantitative ROI-level analysis demonstrated heterogeneous modulation across anatomically defined regions (Fig. S4). Effect-size heatmap visualization revealed distributed changes across frontal, amygdalar, and hippocampal networks, with both increases and decreases in activity (Fig. 4K). Using exact two-sided permutation testing (n = 3 per group), the smallest attainable p-value was 0.10, and no ROI survived FDR correction (all FDR-adjusted p = 0.83). The largest effect was observed in ventral medial entorhinal cortex (ENTm-v), which showed reduced cFos expression in stimulated animals relative to sham (Δ = −2.17; Cohen’s d = −4.75; 95% bootstrap CI: −13.06 to −4.38; Fig. 4L). Within the amygdala, medial amygdala (MeA) demonstrated increased activity under stimulation (Δ = +2.72; d = +1.02; 95% bootstrap CI: +0.88 to +16.39; Fig. 4M), while Anterior cingulate cortex-secondary motor cortex border (ACC–M2) exhibited reduced activation (Δ = −2.61; d = −1.06; 95% bootstrap CI: −45.76 to −0.95; Fig. 4N). Notably, nucleus tractus solitarii (NTS) showed increased activity under stimulation (Δ = +5.00; d = 0.93; bootstrap 95% CI: −0.77 to 4.08; Fig. 4O).

## DISCUSSION

We report a novel, untethered, fully implantable GES for rats to investigate how acute and chronic gastric stimulation modulates anxiety-like behavior in freely moving rats. Unlike previous approaches requiring external tethers or acute surgeries, this device operates continuously in free-moving animals (6, 13, 14, 29), this technological advance allowed us to probe, for the first time to the authors’ knowledge, how stomach stimulation affects anxiety-like behavior over both short and prolonged timescales. Our findings demonstrate that GES exerts context-dependent, temporally dynamic, and phenotype-specific effects on anxiety-related behavior, revealing a level of behavioral organization that is not captured by single measures alone. Critically, the ability to perform chronic stimulation in untethered animals enabled us to dissociate immediate stimulation effects from post-stimulation adaptations, providing new insight into gut–brain mechanisms underlying emotional regulation.

At the level of individual behavioral measures, acute gastric stimulation elicited a robust anxiogenic response. Acute GES was associated with reduced exploratory activity and increased freezing-related behaviors, particularly during stimulation. Freezing is a canonical defensive response in rodents (31), increased freezing coupled with decreased exploratory activity is widely interpreted as heightened anxiety (31). These effects were most pronounced in the OFM, where animals exposed to acute 30 minutes of stimulation displayed elevated anxiety-like behavior relative to controls. In contrast, no significant group differences were detected in the EPM. This dissociation suggests that gastric stimulation does not induce a generalized anxiogenic state, but rather selectively affects contextual and exploratory components of anxiety. This pattern aligns with extensive evidence that the OFM and EPM, although both widely used as assays of anxiety-like behavior, rely on overlapping but partly dissociable neural substrates (6, 34–36); OFM behavior is strongly shaped by striatal, cerebellar, and parietal/entorhinal contributions to exploration, locomotor patterning, and habituation (37–39). In contrast, EPM performance more prominently recruits the ventral hippocampus–medial prefrontal cortex pathway, thalamic reuniens, and ventral hippocampal interneurons that support approach–avoidance decisions under explicit threat (40–43). The selective OFM sensitivity observed here therefore suggests that acute gastric stimulation primarily modulates context-dependent anxiety and exploratory drive, rather than innate avoidance of open or elevated environments.

Whole-brain c-Fos mapping in mice following acute gastric stimulation provides convergent circuit-level support for this interpretation. Anatomically structured and biologically coherent effect-size patterns emerged. The largest raw effect was observed in ventral medial entorhinal cortex (ENTm-v), which showed a marked reduction in activity in stimulated group, suggesting suppression of entorhinal-driven contextual processing. Within the amygdala, multiple nuclei—including central (CeA), basomedial (BMA), medial (MeA), and striatum-like amygdalar territories—exhibited moderate-to-large positive effect sizes, consistent with enhanced limbic recruitment during gastric stimulation. Hippocampal subfields (CA1–CA3, dentate gyrus, subiculum) displayed heterogeneous directional effects, indicating reorganization rather than uniform excitation or suppression. Furthermore, the Nucleus tractus solitarius (NTS) has displayed an increase in activity. Importantly, these distributed trends map onto circuits implicated in contextual anxiety and exploratory control, aligning with the behavioral dissociation observed between OFM and EPM performance.

A key strength of the present study is the distinction between during-stimulation and post-stimulation behavioral epochs. Correlations spanning these epochs revealed that behavioral state during stimulation was associated with behavior after stimulation, suggesting that GES engaged behavioral states whose influence persisted beyond the immediate stimulation window, consistent with state-dependent plasticity observed in non-invasive neuromodulation for anxiety, where multi-session transcranial stimulation produce enduring (>24 h to days) reductions in anxiety and network-level changes rather than transient locomotor suppression or fatigue (44–47). This temporal dimension becomes especially informative when comparing acute and chronic stimulation protocols. During stimulation, acute GES produced clear anxiogenic effects, reflected in velocity, grooming and freezing behaviors. In the post-stimulation period, however, the two groups diverged significantly. Acute GES group, assessed approximately 10 minutes after stimulation offset, sustained elevated anxiety-like behavior, indicating that the anxiogenic effects of acute GES persist beyond immediate stimulation. In contrast, chronic GES group, assessed several hours after the final stimulation session, occupied the opposite end of the distribution in locomotor activity and exploratory behavior. These animals displayed enhanced exploratory behavior, in some measures exceeding that of control animals, although this effect did not reach statistical significance due to the limited sample size. This divergence suggests that while acute gastric stimulation acts as an interoceptive stressor, repeated or prolonged stimulation may engage adaptive mechanisms that bias behavior toward exploration once stimulation ceases. We found timing to be critical: acute GES group post-stimulation measurements captured only the immediate aftereffect of acute stimulation, whereas chronic GES group measurements occurred after a longer recovery period, allowing for network-level plasticity. Such bidirectional, time-dependent effects are well documented in neuromodulation research; for example, acute VNS preferentially engages arousal and stress circuits without clear mood benefit, whereas repeated or chronic VNS produces robust anxiolytic and antidepressant-like outcomes in rodents and progressive clinical improvement over months in humans (48–52).

To capture the integrated structure of anxiety-like behavior, we employed principal component analysis across multiple behavioral variables in both tasks. In both the OFM and EPM, the first principal component reflected a coherent behavioral pattern characterized by increased freezing and reduced exploration, consistent with prior dimensional approaches to rodent anxiety (53–55). Projection onto AnxietyIndex revealed that acute GES animals scored highest on this latent anxiety dimension in the OFM, with post-hoc analyses confirming a significant difference relative to controls. Importantly, the absence of significant group effects on the EPM-derived AnxietyIndex does not imply task independence. Rather, it suggests differential sensitivity of the underlying neural systems to gastric stimulation. The entorhinal trend observed in our c-Fos analysis provides a plausible circuit correlate for this dissociation, as ventral entorhinal cortex primarily contributes to contextual encoding and goal-directed navigation or exploratory navigation (56–58), while influencing threat and fear behaviors mainly through its contextual inputs to hippocampal and amygdala circuits (59, 60). Suppression of entorhinal output could bias animals toward reduced exploratory engagement in the OFM without necessarily altering elevated-arm avoidance in the EPM. Within each maze, exploratory variables covaried coherently; velocity with grooming and center entry in the OFM, and head dipping with open-arm exploration in the EPM, indicating internally consistent behavioral states rather than nonspecific motor effects. Moreover, cross-maze correlations (e.g., OFM velocity with EPM head dipping) mirror previous reports showing that OFM and EPM exploration share an overlapping anxiety-related substrate despite task-specific demands as mentioned above (53). Further supporting the argument that gastric stimulation appears to perturb a shared anxiety-related system, but its behavioral expression depends on environmental context. Hierarchical clustering analyses further demonstrate that gastric stimulation reorganizes anxiety-related behavior at the phenotypic level. Across all pairwise group comparisons, dendrogram similarity metrics were near zero, indicating negligible overlap in clustering structure between groups. Thus, demonstrating that stimulation protocols do not merely shift animals along a shared anxiety continuum, but instead produce qualitatively distinct behavioral organizations.

Mechanistically, these effects are consistent with evidence that gastric inputs engage gut–brain vagal pathways conveying interoceptive signals from the stomach to the nucleus tractus solitarius (NTS), which in turn modulate locus coeruleus (LC) noradrenergic neurons and broader limbic–forebrain circuits involved in affect and motivation (61–64). Gut-innervating vagal afferents have been shown to critically tune anxiety-like behavior via projections from NTS to the central amygdala and related forebrain regions, and vagal hypersensitivity in functional dyspepsia or colitis models induces persistent changes in amygdalar stress-related signaling together with heightened anxiety- and pain-related behavior, which are reversed by vagotomy or dampening peripheral inflammation (65–67). The NTS and amygdalar activation trends observed here are consistent with recruitment of NTS–amygdala pathways during acute gastric stimulation, potentially mediating the transient anxiogenic phenotype. At the same time, heterogeneous hippocampal effects suggest that stimulation engages memory- and context-processing circuits in a more nuanced manner, possibly contributing to the time-dependent behavioral divergence between acute and chronic stimulation. Acute recruitment of these vagal circuits can transiently enhance arousal and stress-related neuromodulation through NTS–LC and NTS–amygdala pathways, contributing to anxiety-related behavioral states in response to gastrointestinal perturbation (61, 62, 65, 66, 68). By contrast, repeated vagal activation engages experience-dependent plasticity in prefrontal–limbic and hippocampal networks and associated memory systems, including anterior cingulate cortex and hippocampus, supporting longer-term changes in anxiety regulation, exploratory behavior, and cognitive control (69–71). The preferential modulation of open-field exploration is consistent with work showing that vagal gut–brain signaling and related neuromodulatory interventions bias activity within hippocampal–prefrontal and striatal-connected circuits that encode contextual anxiety, avoidance of exposed spaces, and the flexibility of exploration–avoidance trade-offs (3, 67, 68, 70, 71).

The present whole-brain mapping therefore serves as a circuit-level foundation upon which future mechanistic studies can build. Increasing sample size will allow formal statistical resolution of the amygdalar and hippocampal trends observed here, and combining chronic stimulation paradigms with longitudinal activity mapping could clarify whether the shift toward exploratory behavior is accompanied by progressive rebalancing of entorhinal–hippocampal and prefrontal–amygdalar networks. Integration with wireless electrophysiology, calcium imaging, or projection-specific manipulations would further delineate how gastric afferent signals reshape limbic circuit dynamics over time. This study tested a FDA-approved GES protocol, and it will be important to vary stimulus parameters and durations to determine differential efficacy in both health and neuropsychiatric disease models. The results described here are consistent with VNS literature.

While the present study focused on healthy male animals and a single stimulation parameter set, future work can expand this framework to stress models, pathological anxiety states, sex differences, and developmental time points to determine how gut–brain neuromodulation interacts with vulnerability or resilience factors. Expanding both behavioral and circuit-level sample sizes will enable more definitive statistical inference while preserving the systems-level approach demonstrated here.

Our findings demonstrate that gastric electrical stimulation reorganizes anxiety-related behavior in a context-dependent, time-dependent, and phenotype-specific manner. Acute stimulation transiently enhances anxiety-like behavior in novel environments, whereas chronic stimulation appears to promote adaptive exploratory behavior following stimulation cessation. Whole-brain c-Fos mapping provides convergent circuit-level support for these behavioral effects, revealing anatomically distributed modulation of entorhinal, amygdalar, NTS and hippocampal networks consistent with altered contextual anxiety processing. By integrating chronic, untethered neuromodulation, brain activity mapping, and multidimensional behavioral analysis, we provide new insight into the gut–brain regulation of anxiety and establishes a powerful framework linking behavior to distributed circuit dynamics.

## METHODS

### Animals

All experimental procedures were approved by the Institutional Animal Care and Use Committee of New York University. All animal care and experimentation were conducted in accordance with the NIH Guide for the Care and Use of Laboratory Animals and institutional ethical guidelines. Every effort was made to minimize animal suffering and to use the minimum number of animals necessary to achieve statistical validity. Twenty-three male Sprague–Dawley rats (body weight 501–550 g) were used in this study. Animals were obtained from Charles River Laboratories and housed in a temperature-controlled vivarium on a 12:12 h light/dark cycle (lights on at 6:30 a.m.). Rats were allowed to acclimate for 3–5 days prior to any experimental procedures, with food and water available *ad libitum*. Prior to surgery, animals were pair-housed; after surgical implantation they were housed individually to prevent interference with externalized leads. To anatomically resolve central activation patterns induced by gastric electrical stimulation, an independent cohort of adult male C57BL/6 mice (n = 6; 8–10 weeks old; 20–30 g) was used for whole-brain c-Fos mapping. Mouse experiments were performed separately from the rat behavioral experiments and were not pooled for statistical analysis. Stimulation parameters were matched to those used in rats (14 Hz, 0.3 ms pulse width, 0.5 mA) to ensure cross-species comparability. Mice were housed under identical light/dark conditions with ad libitum access to food and water.

### Surgical procedures

All surgeries were performed under aseptic conditions, by one experimenter, in 3 consecutive days, and isoflurane anesthesia delivered via a VetEquip vaporizer (VetEquip Inc., Pleasanton, CA). Anesthesia was induced at 3% isoflurane and maintained at ~2.5%, with adjustments made as needed according to respiratory rate. Meloxicam (1 mg/kg, covetrus, UK) and eye ointment (Muro, Bausch+Lomb, USA) were administered. The rat was placed supine on a heating pad (CVS heating pad, USA) to maintain body temperature. For electrode implantation (see Fig.S1D for step by step pictures), a midline laparotomy was performed to expose the stomach. Two 14.5-inch platinum–iridium wires (0.003” diameter) were implanted into the mid-to-lower portion of the greater curvature of the stomach, approximately 1 cm apart. Only 0.5 cm of the electrodes were laser cut (Epilog Zing 24 Laser Cutter) to remove the coating and this part was in contact with the mucosa of the stomach (Fig. S1C). A short segment of flexible polymer tubing (inner diameter ~0.011”) was placed over each electrode tip to increase rigidity, and the electrode tips with tubing were secured to the serosal surface of the stomach using a few drops of surgical tissue adhesive. An additional segment of polyethylene tubing (inner diameter ~0.045”) was placed over each pair of electrode tips for strain relief. The electrode leads were then routed subcutaneously to the dorsal side using a sterile trocar and externalized through a small incision just above the lumbar vertebrae. Each lead was soldered to a connector terminal. The connector assemblies were secured in a custom dorsal backpack housing (see below). The abdominal muscle layer and skin layer were closed separately with 5-0 VetLON sutures (simple interrupted pattern; Veter.Sut FS-2). An additional Meloxicam dose (1 mg/kg) and 0.9% saline (5 mL/kg) were administered at the end of surgery. Surgeries were done under an hour total time. Post-operative care included analgesia: rats received meloxicam (2 mg/kg) once daily for 3 days. They were also provided with hydration gel. Animals were closely monitored during a 3-day recovery period before behavioral testing commenced. Experimental groups and sham rats were submitted to the same procedures. For the mice cohort, a lateral abdominal incision was made inferior to the left hypochondriac region. Following laparotomy, bipolar needle electrodes were inserted trans-serosally through the stomach wall mid-to-lower portion of the greater curvature, 2 mm apart. Electrical stimulation was applied for 15 minutes. Sham-operated mice underwent electrode implantation without stimulation. Animals were maintained under deep anesthesia for 45 minutes post-stimulation to allow c-Fos expression, then immediately euthanized via transcardial perfusion (PBS + heparin) followed by 4% paraformaldehyde (PFA). Brains were dissected, post-fixed in 4% PFA overnight at 4°C, and stored in 1× PBS with 0.02% sodium azide at 4°C until processing.

### GES backpack and stimulation circuit

GES was delivered by a custom-designed stimulator in a dorsal-mounted backpack. The total backpack unit weighed approximately 30 g and housed a custom printed circuit board (PCB) and a rechargeable lithium-polymer battery, enabling fully untethered operation during behavioral testing. The dorsal mounting preserved natural posture and locomotion. Stimulation parameters were fixed at 14 Hz with 0.3 ms monophasic pulses (0.5 mA amplitude), parameters that are used clinically in GES (30). These parameters were set via adjustable potentiometers and swappable timing capacitors on the PCB, allowing precise and reproducible configuration across devices while maintain identical stimulation conditions across animals. The stimulation architecturally consists of a 555 timer feeding into a constant current source integrated circuit. Output from the constant current source is robust to change in tissue impedance that may arise from posture, movement or gastric motility. Each fully charged device could provide more than 48 hours of continuous stimulation, enabling uninterrupted stimulation across behavioral testing sessions without battery replacement or mid-experiment intervention. Proper function and pulse delivery were tested on a benchtop physiologically representative load via an oscilloscope (SigLent SDS 1104X-E) prior to in vivo use. The backpack was placed on a strong Velcro on top of a rat jacket. We have custom tailored and padded the rats’ jacket from Lomir Biomedical (Fig. S1F) to prevent irritations under armpit. The padding has successfully prevented irritations and ulcers post 14 days (see Fig. S1G).

### Behavioral testing

Tests were conducted during the light circle. These tests were chosen because they utilize the rats’ innate aversion to open, brightly lit, or elevated spaces to assess anxiety responses (31).

#### Open Field Maze (OFM)

The OFM apparatus was a square open-field arena measuring 100 cm x 100 cm with 35 cm high opaque walls, obtained from San Diego Instruments (San Diego, CA). A central zone (50 cm x 50 cm) was defined within the arena to evaluate center vs. periphery exploration, as rodents avoid the open center, increased thigmotaxis is interpreted as higher anxiety. The OFM was also used to measure locomotion behavior such as velocity, defecation, and latency to enter the center.

#### The Elevated Plus Maze (EPM)

The EPM apparatus consisted of two opposing open arms (114 cm x 10 cm) and two opposing closed arms (114 cm x 10 cm, with 40 cm high walls) arranged in a plus configuration and elevated 50 cm above the floor, obtained from San Diego Instruments (San Diego, CA). The percentage of time spent in the open and enclosed arms as a function of total arm time and the rate of entries into the open and enclosed arms as a function of all arm entries are measured. Increased time and entries into the enclosed arms are interpreted as greater avoidance of elevated, open spaces.

Both apparatuses were cleaned with 70% ethanol between rats to eliminate olfactory cues. Behavioral testing took place on specific days post-surgery, depending on group assignment (see Stimulation Protocol below for schedule). Each rat was gently placed in the center of the apparatus at the start of each session and allowed to freely explore. Sessions were video recorded from above for subsequent behavioral analysis.

### Stimulation protocol

We used four experimental groups to dissociate acute vs. chronic effects of GES, with or without surgery, as follows (see s. S1A):

#### Group 1 (Acute GES)

This group received short-term gastric stimulation only during behavioral testing sessions. In each test session, GES was delivered for 30 min while the rat was in the apparatus. The stimulation period was preceded by a 10-min baseline with the stimulator off and followed by a 10-min post-stimulation period with the stimulator off (total session time 50 min). Group 1 underwent three such test sessions (for each open field and elevated plus maze) over the 2-week experimental period (on post-surgery days 5–6, 8–9, and 11–12, with rest days between) to examine within-subject acute effects of GES.

#### Group 2 (Chronic GES)

This group received extended, continuous GES to model chronic stimulation. Each Group 2 rat underwent two 48-hour stimulation periods: one spanning days 5–6 post-surgery and another spanning days 9–10 post-surgery. Behavioral tests were administered during, after 48 h stimulation and after 48 h of no stimulation period to assess the effects of sustained GES. In practice, animals had behavioral testing at roughly the midpoint-end of the 48 h stimulation, immediately after the second 48 h stimulation, and follow-up testing after the stimulation was turned off for 48 h. The stimulator’s battery was recharged between these two stimulation periods. This chronic protocol enabled evaluation of both within-session and longer-term (days apart) effects of GES.

For both experimental groups, sweep impedance measurements were taken before each stimulation period to ensure integrity of electrodes, while the rats were awake and freely moving. Impedance measurements were taken by connecting the circuit board to an oscilloscope, concurrently rats were handled and calmed to stay still while the measurements were completed.

#### Group 3 (Sham Surgery)

This group served as a sham stimulation control. Group 3 animals underwent the same surgical procedure as the GES groups (electrodes implanted and connected to the backpack), but the stimulator was never activated. These rats were handled and tested on the same schedule as the chronic GES group, including being outfitted with the backpack during test sessions, but they received no electrical stimulation. This allowed us to control for the effects of surgery and the presence of the device without the confounding influence of GES current.

#### Group 4 (No Surgery Control)

This group consisted of rats that did not undergo any surgery or electrode implantation. To match the handling and environmental exposure of the other groups, Group 4 rats were habituated to wearing a dummy backpack, of similar weight to the real stimulator but containing no electronics, during the experiment. They were handled daily and placed into the testing apparatus on the same days as the other groups. This no-surgery group controlled for any effects of surgery on behavioral measures.

### Stimulation paradigm for c-Fos induction

In mice, a single acute stimulation (14 Hz, 0.3 ms, 0.5 mA) was delivered for 15 minutes under anesthesia via a TDT stimulator (Tucker Davis Technologies). Sham mice underwent electrode placement without stimulation. Following stimulation, mice remained under deep anesthesia for 45 minutes to allow c-Fos expression prior to transcardial perfusion.

### Automated pose estimation and behavioral quantification

Each behavioral session was recorded using a fixed, top-down camera setup at 40 frames per second with a spatial resolution of 1080 × 1080 pixels, centered on the behavioral maze. We used our in-house camera system which consisted of IMX477 12.3-megapixel CMOS image sensor and Computar 8mm lens to minimize geometric distortion (fisheye problem) and get accurate position of rats in the mazes. Camera system was supported by Raspberry Pi. Setup was fixed throughout the entire experiment to ensure consistent spatial geometry across session. Spatial calibration was performed using known maze dimensions to derive single pixel to centimeter conversion factor, which was applied across all the session uniformly.

Video recordings of each behavioral session were analyzed offline using a custom machine-learning-based tracking pipeline developed in-house. We used a deeplabcut (25) based pose estimation pipeline with a Resnet-152 backbone pretrained on ImageNet. (32). The network was fine-tuned on an in-house curated dataset of rat videos with human annotated keypoints 600 frames collected from different rodents and different sessions, learning to identify and track specific points on the rat’s body. We tracked 16 keypoints including snout, all 4 paws, tail, ears and edges of box on each animal (see Fig. S1B) with a mean tracking error of 12 pixel (out of 16 keypoints 2812×2812pixels) on a held-out test set. The trained model was used to process each frame to generate time series of each keypoint’s 2D coordinates. From these coordinate data, the animal’s moment-to-moment location, orientation, and posture were reconstructed for subsequent behavioral analysis.

Using the extracted pose and trajectory data, we quantified various anxiety-related behaviors. Freezing (immobility) was operationally defined as periods during which the animal’s instantaneous velocity fell below a very low threshold (e.g., < 10 pixels movement) for a sustained duration of at least 0.5 seconds. Grooming bouts were identified based on characteristic repetitive movements of the head and forepaws towards the body (often accompanied by little locomotion), detected either by specific pose configurations or through manual scoring of video if needed, grooming episode was identified when the two fore paws were close to the snout. Rearing events (vertical exploration) were counted whenever the rat stood on its hind legs, which was inferred from an extended vertical displacement of the upper body keypoints or when the forelimb keypoints were elevated above a threshold height. Maze entries were automatically scored by evaluating the rat’s center-point trajectory relative to predefined zone boundaries in the apparatus: for the OFM, an entry into the center zone was counted each time the rat’s center-point moved from the peripheral zone into the 50×50 cm center area (with all four paws inside); for the EPM, open-arm and closed-arm entries were counted when the center-point (and all four paws) crossed from the central platform into an arm. Locomotor movement velocity was computed as the displacement of the rat’s center-point between consecutive video frames, multiplied by the frame rate to yield cm/s. To reduce high-frequency noise, the velocity time series was smoothed with a Butterworth filter and pooling avg distance traveled every 0.25 sec (10 frames). Lastly, defecation rate was manually tracked by the experimenter. These quantitative measures of behavior were used for subsequent statistical analysis of anxiety-like outcomes.

## Statistical Analysis

All behavioral data were reported as the mean ± SEM, and statistical significance was set at p < 0.05 (two-tailed). Where applicable, effect sizes (Cohen’s *d* or partial η^2^) are reported for significant comparisons. Data were analyzed using one-way ANOVA (data from OFM and EPM separately for pre-stim, during stim and post stim and *Group* as between-subject factor) and one-way repeated measures ANOVA (with *Epoch* as within-subject factor), followed by Tukey’s multiple comparison test. When using repeated measures ANOVA, Mauchly’s Sphericity Test was performed, and corrections were made using Greenhouse-Geisser (G-G). Additionally, for Group 1 (Acute GES), within-session comparisons were made across the different stimulation phases. Each test session for Group 1 was divided into five consecutive epochs: a pre-stimulation baseline, three 10-min stimulation blocks (covering the 30 min of GES ON), and a post-stimulation epoch. Linear mixed-effects models were used to test differences across these epochs (within-subject factor: *Epoch*). Post-hoc comparisons between specific epochs were Bonferroni-corrected where applicable. Furthermore, for Group 2 (Chronic GES), longitudinal changes across testing days were analyzed with linear mixed-effects model. The within-subject factor was *session* (i.e. during stimulation, post stimulation and wash off). Throughout the study, a total of five animals did not complete the entire protocol: three rats were lost following an accidental insufficient-oxygen event, and in two additional rats the jacket became caught on a tooth, prompting Division of Comparative Medicine (DCM) staff to remove the backpack entirely to prevent injury; data from these animals were included up to the last successfully completed session.

In addition to ANOVA/GLM analyses, we explored relationships between different behavioral measures (i.e. OFM and EPM) across different epochs (i.e. during stimulation and post stimulation). Missing values were handled using multiple imputation in SPSS (five imputations), and the pooled dataset was used for all subsequent analyses to reduce bias associated with listwise deletion. Pearson correlation coefficients were computed between all behavioral variables. To control for type I error due to multiple testing, p-values were adjusted using the Benjamini–Hochberg False Discovery Rate (FDR) procedure. We also probed potential moderation effects to see if the impact of GES on behavior was dependent on *Group* (Acute GES, Chronic GES, Sham Surgery, Control). For these moderation analyses, we used ordinary least squares (OLS) regression models in which an interaction term between *Epoch* and *Group* (with group effect coded as a four-level dummy variable) was included to assess if group significantly altered the relationship between a predictor and an outcome.

To capture a multivariate anxiety-like behavior dimension, we performed principal component analysis (PCA) separately on the OFM and EPM data (each pooled across imputations). All behavioral variables were z-scored prior to PCA. For the OFM, variables included average speed, center zone time, center entries, periphery time, freezing time, grooming time, rearing time, freezing rate, grooming rate, rearing rate, and defecation count. For the EPM, variables included percent time on open arms, percent entries into open arms, freezing rate, an open-arm exploration index (calculated as 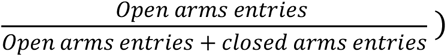, head-dip rate, stretch-attend posture (SAP) rate, freezing time, and defecation rate. In both tasks, the first principal component (PC1) captured the canonical anxiety-related pattern (high freezing and low exploration). We multiplied PC1 by–1 so that higher values consistently indicate greater anxiety-like behavior. This unified “AnxietyIndex” was used as the primary outcome variable in all models, though key individual measures (e.g. locomotion, center exploration, open-arm time) were also examined in parallel to interpret the multivariate results. Group and epoch effects on anxiety-related behavior were tested using linear mixed-effects models (LME) in SPSS (REML estimation, Satterthwaite correction for degrees of freedom). The main model took the form: AnxietyIndex ~ Group × Epoch + (1|Rat_ID), where Group has four levels (Acute GES, Chronic GES, Sham Surgery, No-Surgery Control) and Epoch has up to three levels depending on group (for Group 1 [Acute GES]: Pre, During, Post; for Groups 2–4: During, Post). In these models, Rat_ID was treated as a random intercept to account for repeated measures on each animal. Significant main effects or interactions were followed by Tukey-corrected pairwise contrasts (two-tailed).

Furthermore, to examine whether the multivariate structure of behavioral variables differed between stimulation groups, hierarchical clustering was performed at the variable level within each group. All variables were standardized (z-scored) within group, and pairwise Euclidean distances between variables were calculated using pairwise-complete observations. Distance matrices were embedded into Euclidean space via classical multidimensional scaling (MDS) to satisfy the assumptions of Ward’s minimum variance method. Hierarchical clustering was then conducted using Ward’s linkage with optimal leaf ordering. The optimal number of clusters (k) was determined by inspecting the scree plot (elbow method) of the agglomeration distances. Structural similarity between groups was further quantified by comparing dendrograms pairwise using cophenetic coefficients and Baker’s Gamma rank correlation.

We also analyzed electrode impedance (|Z|) spectra to compare tissue/electrode interface properties across conditions. Impedance magnitude was measured across a range of frequencies for each implanted electrode [10 Hz - 1,000,000 Hz]. For each subject, the raw impedance vs. frequency data were linearly interpolated onto a common frequency grid to allow averaging. We then averaged the impedance spectra within each group (Acute GES, Chronic GES, Sham Surgery) and compared these using a cluster-based permutation testing approach. Specifically, at each frequency point we computed Welch’s t-test between two groups of interest. Clusters of consecutive frequency points showing t-test *p*-values below a threshold (i.e. *p* < 0.05) were identified, and a null distribution of cluster sizes was obtained by performing 5,000 random permutations of group labels. A cluster was considered significant if its size (frequency span) exceeded the 95th percentile of the null distribution.

All statistical analyses were performed using IBM SPSS Statistics (version 31) and GraphPad Prism (version 10) and Python (version 3.0).

### Histology and tissue analysis

At the conclusion of the experiment (day 14 post-surgery), all rats who underwent surgery were euthanized for tissue analysis. Rats were deeply anesthetized with isoflurane vapor (5% in oxygen) and then overdosed. While under deep anesthesia, they were transcardially perfused with 0.9% saline followed by 4% paraformaldehyde in phosphate buffer to fix the tissues. The stomach, particularly the region of electrode implantation, was then carefully harvested. Tissue samples were put into a solution of 10% formalin for 72 hours, transferred into a 70% ethanol solution, and submitted to the NYU Histology Core for processing. Serial sections of the stomach wall encompassing the electrode tract were mounted on slides and stained with hematoxylin and eosin (H&E) following standard protocols to examine tissue morphology. A board-certified pathologist examined the samples.

### Tissue Clearing and Imaging

For iDISCO+ processing, mouse brains underwent dehydration in graded methanol/H_2_O solutions (20-100%), followed by overnight incubation in 66% dichloromethane/33% methanol. After methanol washes and overnight bleaching in 5% H_2_O_2_/methanol at 4°C, samples were rehydrated through graded methanol/H_2_O solutions and PBS. Permeabilization was performed for 2 days at 37°C, followed by blocking for 2 days at 37°C. Primary antibody incubation used rabbit anti-c-Fos (Synaptic Systems, #226 003, 1:2000) for 7 days at 37°C. Following washes, secondary antibody incubation (donkey anti-rabbit Alexa Fluor 647, 1:500) was performed for 7 days at 37°C. Final clearing involved re-dehydration, dichloromethane treatment, and refractive index matching in dibenzyl ether for ≥24 hours before imaging. For CLARITY processing, brains were incubated in A4P0 hydrogel solution for 24 hours at 4°C, degassed, and polymerized at 37°C for 2 hours. Clearing was performed using the X-CLARITY system with 4% SDS solution. After washing, samples were incubated in RIMS solution (refractive index 1.45-1.485) for ≥24 hours before imaging. Imaging was performed using a CLARITY-optimized light-sheet microscope with pixel resolution of 1.4 × 1.4 µm^2^ (x-y dimensions) and 5 µm z-step using a 4×/0.28 NA objective. Two channels were used: autofluorescence (488 nm) for brain anatomy and 647 nm for c-Fos labeling. Raw 16-bit TIFF images were stitched using TeraStitcher

### Whole-brain atlas registration and cell quantification

Whole cleared brains were registered to the Allen Mouse Brain Atlas and analyzed using the AI-based Cartography of Ensembles (ACE) pipeline within MIRACL (Multimodal Image Registration And Connectivity anaLysis) (33). Briefly, c-Fos expression was segmented using deep learning-based vision transformer architectures (UNET/UNETR models) trained on large-scale light-sheet fluorescence microscopy datasets. Segmented c-Fos-positive neurons were voxelized and warped to atlas space for automated anatomical assignment. Cluster-level anatomical annotations derived from the Allen Brain Atlas were normalized by removing cortical layer identifiers and mapped to granular anatomical regions of interest (ROIs) using an ordered, rule-based dictionary with predefined priority resolution to ensure deterministic assignment of each cluster to a single ROI. White-matter and ventricular annotations were excluded from assignment unless they were the sole label. This procedure yielded granular ROIs spanning medial prefrontal cortex, anterior cingulate and motor cortex, amygdala subdivisions, hippocampal subfields, and entorhinal cortex. For each animal, ROI-level cFos values were computed as the mean across all clusters assigned to that ROI. Group differences (stimulation vs sham) were assessed independently for each ROI using exact two-sided permutation tests by enumerating all possible label assignments (n = 3 per group). Effect sizes were quantified as Cohen’s d (difference in group means normalized by pooled standard deviation). Uncertainty in effect sizes was estimated using a nonparametric bootstrap within each group to obtain percentile-based 95% confidence intervals for Cohen’s d. Multiple comparisons across ROIs were controlled using the Benjamini–Hochberg false discovery rate (FDR; α = 0.05). Given the small sample size, statistical analyses were treated as exploratory. Analyses were automated and reproducible using a custom analysis script

## Acknowledgments

We thank the NYU Experimental Pathology Research Laboratory (RRID:SCR_017928) for histological processing and analysis, supported in part by the NYU Cancer Institute Center Support Grant (NIH/NCI P30CA016087). We acknowledge the support of NYU Information Technology High Performance Computing resources, services, and staff expertise, which were essential for behavioral data processing and analysis. We also acknowledge NYU LH Microscopy Lab (RRID: SCR_017934) for providing microscopy services. The microscopy shared resource is partially supported by Cancer Center Support Grant P30CA016087. We thank the Division of Comparative Medicine staff at New York University for their assistance with animal care and surgical oversight. This work was supported by the Advanced Research Projects Agency for Health (ARPA-H).

