## Supplementary material for "Divergent Behavioral and Circuit-Level Adaptations to Acute and Chronic Gastric Electrical Stimulation"

Supplemental materials:

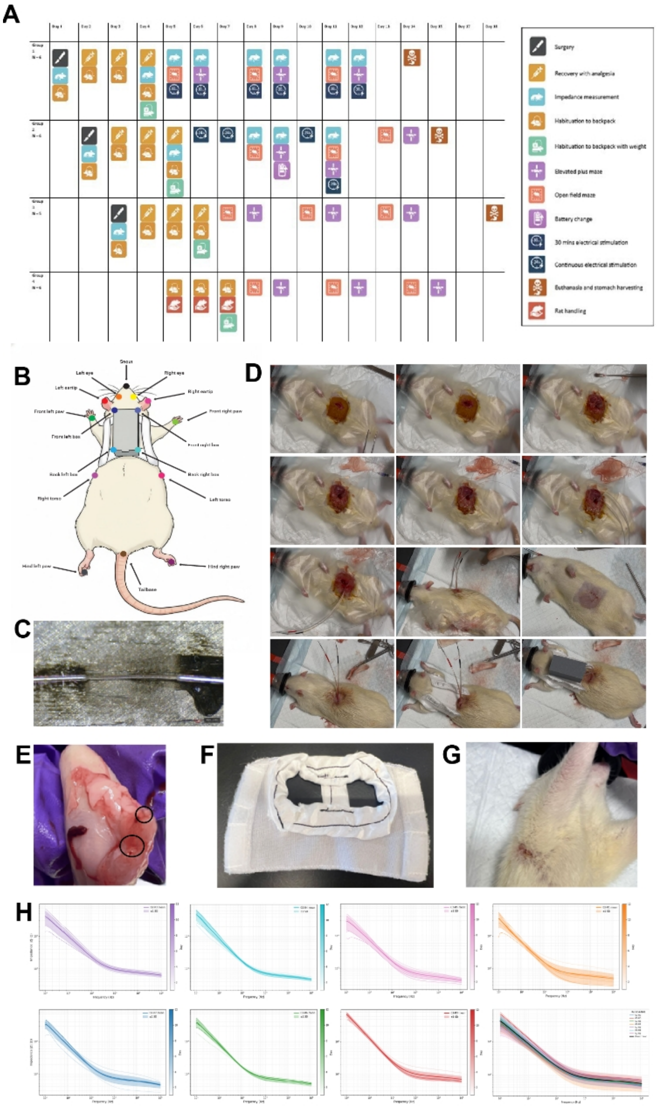


**Figure S1. Experimental timeline, pose estimation, surgical refinement, and device stability.** (A) Experimental timeline illustrating four groups: Acute GES, Chronic GES, Sham Surgery, and

No-Surgery Control. Acute GES animals received 30-minute stimulation during behavioral testing; Chronic GES animals received two 48-hour stimulation blocks; Sham and Control groups underwent identical handling without active stimulation. (B) Keypoint configuration for DeepLabCut pose estimation (16 keypoints). (C) Laser-cut exposed electrode tip. (D) Step-by-step intraoperative implantation images. (E) Postmortem confirmation of electrode placement (n=12). (F) Custom-fitted padded jacket with satin fabric. (G) Skin condition after 14 days of wearing the system. (H) Sample individual impedance trajectories over time (mean ± SD band).

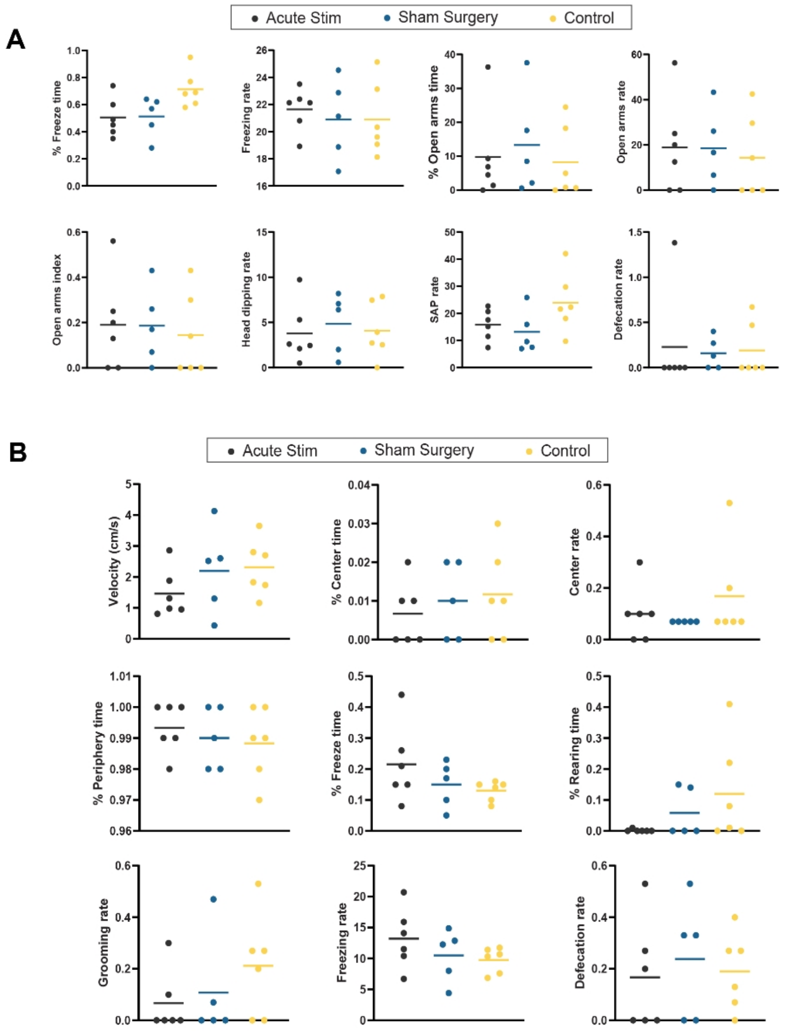


**Figure S2. Baseline behavioral equivalence.** Pre-stimulation OFM and EPM metrics across groups (Acute n = 6; Sham n = 5; Control n = 6). One-way ANOVAs revealed no significant baseline differences among the three groups on any measured behavioral metric.

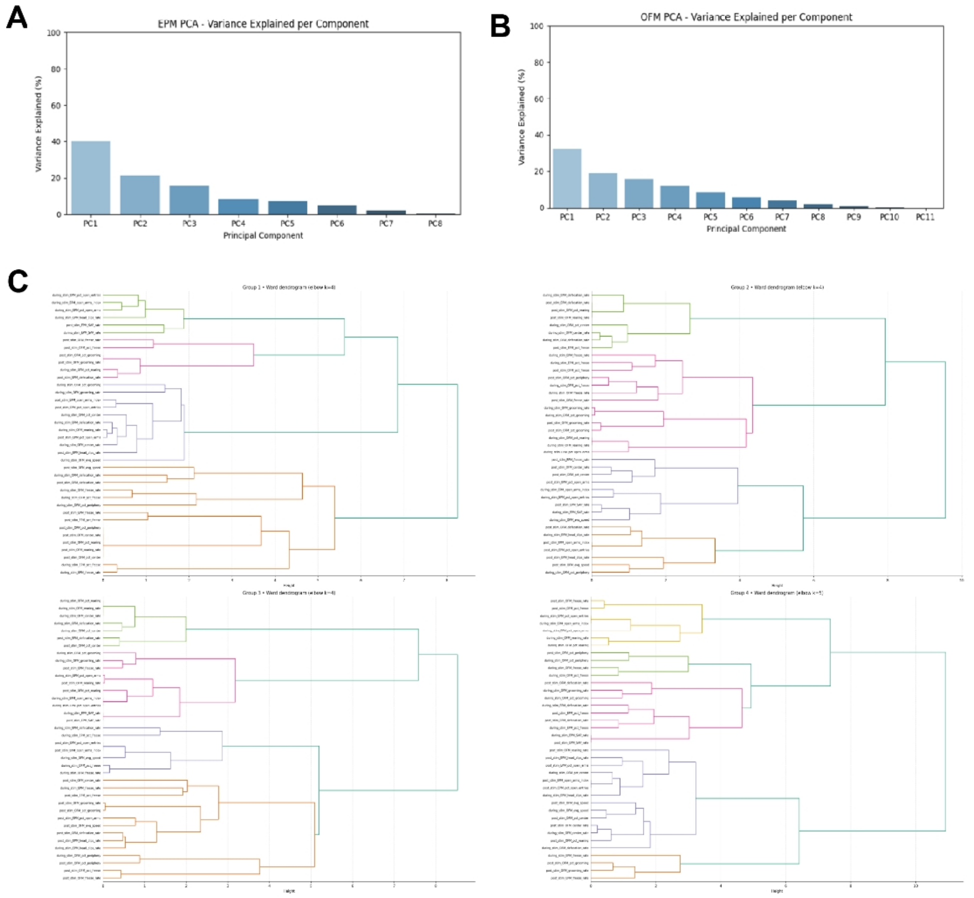


**Figure S3. Principal component analysis and clustering.** (A) Scree plot and loadings for OFM PCA (n = 19 animals; PC1 explains 31.9% variance). (B) Scree plot and loadings for EPM PCA (n = 19 animals; PC1 explains 39.8% variance). (C) Hierarchical clustering dendrograms computed using Ward linkage on z-scored behavioral variables. Across all pairwise group comparisons, cophenetic correlation coefficient (CCC) values were extremely low (0.059–0.157) and Baker's Gamma coefficients were close to zero (−0.008–0.077).

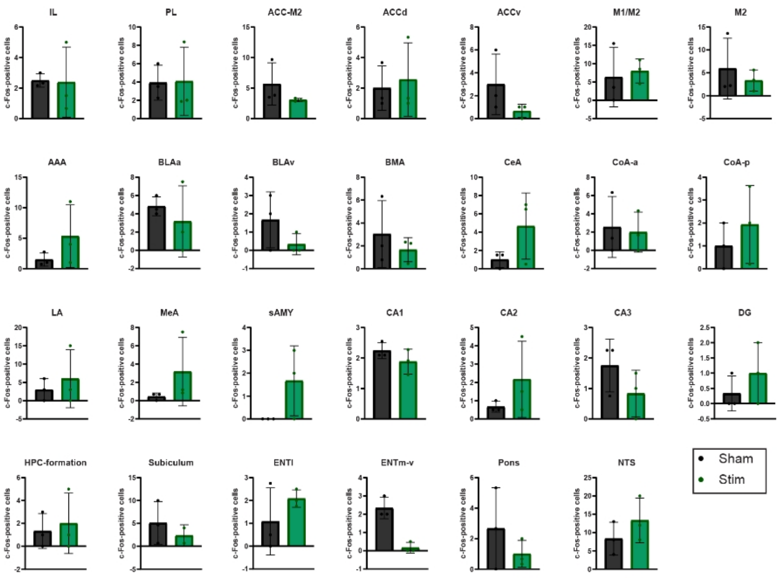


| Region | ROI | Mean Sham | Mean Stim | Delta | p value | FDR value |
|---|---|---|---|---|---|---|
| **Frontal-mPFC** | | | | | | |
| | IL | 2.500 | 2.389 | -0.111 | 1.0000 | 1.0000 |
| | PL | 3.917 | 4.083 | 0.167 | 1.0000 | 1.0000 |
| **ACC/Motor** | | | | | | |
| | ACC-M2 | 5.667 | 3.056 | -2.611 | 0.1000 | 0.8000 |
| | ACCd | 2.000 | 2.556 | 0.556 | 0.9000 | 1.0000 |
| | ACCv | 3.000 | 0.667 | -2.333 | 0.3000 | 0.8000 |
| | M1/M2 | 6.333 | 8.000 | 1.667 | 0.8000 | 1.0000 |
| | M2 | 5.933 | 3.333 | -2.600 | 0.8000 | 1.0000 |
| **Amygdala** | | | | | | |
| | AAA | 1.500 | 5.333 | 3.833 | 0.3000 | 0.8000 |
| | BLAa | 4.833 | 3.167 | -1.667 | 0.5000 | 0.8235 |
| | BLAv | 1.667 | 0.333 | -1.333 | 0.4000 | 0.8000 |
| | BMA | 3.037 | 1.667 | -1.370 | 0.6000 | 0.9333 |
| | CeA | 1.000 | 4.667 | 3.667 | 0.3000 | 0.8000 |
| | CoA-a | 2.556 | 2.000 | -0.556 | 0.9000 | 1.0000 |
| | CoA-p | 1.000 | 1.933 | 0.933 | 0.5000 | 0.8235 |
| | ITC | 0.667 | 0.667 | 0.000 | 1.0000 | 1.0000 |
| | LA | 3.000 | 6.000 | 3.000 | 0.9000 | 1.0000 |
| | MeA | 0.444 | 3.167 | 2.722 | 0.1000 | 0.8000 |
| | sAMY | 0.000 | 1.667 | 1.667 | 0.4000 | 0.8000 |
| **Hippocampus/DG/Sub** | | | | | | |
| | CA1 | 2.242 | 1.879 | -0.364 | 0.4000 | 0.8000 |
| | CA2 | 0.667 | 2.167 | 1.500 | 0.4000 | 0.8000 |
| | CA3 | 1.750 | 0.833 | -0.917 | 0.3000 | 0.8000 |
| | DG | 0.333 | 1.000 | 0.667 | 0.7000 | 1.0000 |
| | HPC-formation | 1.333 | 2.000 | 0.667 | 0.9000 | 1.0000 |
| | Subiculum | 4.889 | 1.778 | -3.111 | 0.4000 | 0.8000 |
| **Parahippocampal (EC)** | | | | | | |
| | ENTI | 1.083 | 2.083 | 1.000 | 0.4000 | 0.8000 |
| | ENTm-v | 2.333 | 0.167 | -2.167 | 0.1000 | 0.8000 |
| **Brainstem** | | | | | | |
| | Pons | 2.667 | 1.000 | -1.667 | 0.4000 | 0.8000 |
| | NTS | 8.333 | 13.333 | 5.000 | 0.5000 | 0.8235 |

**Figure S4. ROI-level statistical from whole-brain c-Fos analysis.** Detailed ROI-level group comparisons, plots, mean normalized cFos density (cells/mm³) per group, delta, effect sizes (Cohen's d), Two-sided permutation p-value ,and FDR-adjusted q-values across anatomically defined regions. Acute GES n = 3; Sham n = 3. FDR correction applied across all ROIs (Benjamini–Hochberg, q = 0.05). ROIs shown are: Infralimbic cortex (IL); Prelimbic cortex (PL); Dorsal anterior cingulate cortex- secondary motor cortex border (ACC-M2); Dorsal anterior cingulate cortex (ACCd); Ventral anterior cingulate cortex (ACCv); Primary/secondary motor cortex (M1-M2); Secondary motor cortex (M2); Anterior amygdalar nucleus (AAA); Anterior basolateral amygdalar nucleus (BLAa); Ventral basolateral amygdalar nucleus (BLAv); Basomedial amygdalar nucleus (BMA); Central amygdalar nucleus (CeA); Anterior cortical amygdalar area (CoA-a); Posterior cortical amygdalar area (CoA-p); Intercalated amygdalar nucleus (ITC); Lateral amygdalar nucleus (LA); Medial amygdalar nucleus (MeA); Striatum-like amygdalar nuclei (sAMY); Hippocampal field CA1; Hippocampal field CA2; Hippocampal field

CA3; Dentate gyrus (DG); Hippocampal formation; Subiculum; Lateral entorhinal cortex; Ventromedial entorhinal cortex; Pons; Nuvleus tractus solitarii (NTS).
